# Role of transcription and translation during the early development of the brown alga *Ectocarpus*

**DOI:** 10.1101/2023.10.19.563035

**Authors:** Daniel Liesner, Rémy Luthringer, Sébastien Colin, Julia Morales, J. Mark Cock, Susana M. Coelho

## Abstract

**Background and aims:** Parthenogenesis, the embryonal development of an unfused gamete, is a widespread trait within the brown algae (Phaeophyceae). We hypothesized that the parthenogenetic development of male gametes of the model brown alga *Ectocarpus* species 7 would rapidly be dependent on *de novo* transcription and translation because of the small size of the gamete cell.

**Methods:** We followed the development of male *Ectocarpus* gametes to parthenosporophytes in the presence of either the transcription inhibitor thiolutin or the translation inhibitor emetine. Responses in morphology and growth were compared to development in inhibitor-free control conditions at three time points over 12 days. Potentially persistent inhibitor effects were then investigated by growing parthenosporophytes in an inhibitor-free post-culture for 14 days.

**Key results:** Thiolutin did not affect gamete germination, but growth of parthenosporophytes was significantly delayed. While almost all control parthenosporophytes had grown larger than 10 cells over 12 days, thiolutin inhibited growth beyond a size of 5-10 cells. The effects of thiolutin were reversible in the post-culture. Consequences of the emetine treatment were more severe, germination was already strongly inhibited by day 5, and on average only 27.5% of emetine-treated gametes had completed the first cell division on day 12. Emetine fully inhibited development beyond the 5-cell stage during the treatment, and induced morphological abnormalities (i.e., round cell shape and abnormal cell division planes) which persisted throughout the post-culture.

**Conclusions:** These results imply that *Ectocarpus* gametes contain sufficient proteins to germinate, and that the first cell cycles of parthenogenetic gamete development presumably utilize mRNA already present in the gametes. We discuss that storing mRNA and proteins in the developing gametes before release may be an adaptive trait in *Ectocarpus* to ensure quick development after fertilization, or alternatively the vegetative completion of the life cycle in the absence of mates.

## Introduction

Brown algae have been used for decades as models to study embryogenesis (Coelho *et al*., 2020; Bogaert *et al*., 2023). This is because they offer a number of advantages such as the ease with which gametes and zygotes can be obtained and manipulated (contrary to land plant systems where the embryos are embedded in parental sporophytic tissue), their suitability for cellular imaging studies and microinjection, coupled with the ability to carry out biochemical analyses of large numbers of synchronously developing zygotes (Brownlee *et al*., 2001; Bogaert *et al*., 2013; Coelho and Cock, 2020). Thorough characterization of polarization, germination and first cell divisions in this group of organisms has revealed that early developmental processes are crucial in the determination of the correct patterning of the embryo and future adult (Robinson and Miller, 1997; Brownlee and Bouget, 1998; Corellou *et al*., 2000, 2005; Pool *et al*., 2004). These studies, however, focused on organisms where a large female gamete (egg) is fertilized by a small male gamete (sperm; *i*.*e*., oogamy), such as in the morphologically most complex brown algal orders of wracks (Fucales) and kelps (Laminariales). While oogamy is considered the ancestral state in the brown algae (Silberfeld *et al*., 2010; Heesch *et al*., 2021), this group actually exhibits an exceptionally broad range of sexual systems, ranging from isogamy to oogamy with different degrees of sexual differentiation (Silberfeld *et al*., 2010; Luthringer *et al*., 2014; Heesch *et al*., 2021). For instance, in many brown algal species male and female gametes have approximately the same size (*i*.*e*., near-isogamy).

Parthenogenesis is a form of asexual reproduction in which a haploid gamete develops embryonically despite the absence of gamete fusion. In the predominantly oogamous plants and animals, parthenogenesis is restricted to the larger gametes, *i*.*e*. the eggs. Similarly, in many oogamous brown algae species, unfertilized eggs can develop into parthenosporophytes (Heesch *et al*., 2021) which may be morphologically identical to sporophytes obtained by gamete fusion (Peters *et al*., 2008; Hoshino *et al*., 2019) or present abnormalities in physiology and morphology presumably due to their haploid genomes (tom Dieck, 1992; Müller *et al*., 2019). The triggering of embryonic development is therefore independent of fertilization and ploidy (Bothwell *et al*., 2010; Coelho *et al*., 2011). Interestingly, (near-)isogamy in brown algae is often associated with the capacity of both gamete sexes to develop parthenogenetically (Luthringer *et al*., 2014). In all cases of parthenogenesis, the early developmental program has to be initiated and sustained in the presence of only one parental genome.

Haploid gametophytes of the filamentous model brown alga *Ectocarpus* produce small (approximately 4 μm) male and female near-isogametes, which, upon gamete fusion, initiate the sporophyte generation (**Figure 1**; Müller, 1967). For most *Ectocarpus* strains, gametes that do not fuse with a partner of the opposite sex are able to develop parthenogenetically in an asexual cycle (Müller, 1966; Bothwell *et al*., 2010). Compared to zygotic development, which is triggered immediately following gamete fusion, parthenogenetic development of unfused gametes proceeds after a delay of at least 24 h and growth is reduced during the first few days of development (Peters *et al*., 2008). Apart from this delay, the pattern of early development of a diploid sporophyte (zygote as initial cell) and a parthenosporophyte (gamete as initial cell) is largely identical (Peters *et al*., 2008). Following gamete settlement, the round initial cell elongates during germination (**Figure 1**). Germination is asynchronous but bipolar, and the two daughter cells produce the two ends of a symmetric prostrate filament which grows by further cell divisions within a few days. The cells of the prostrate filament become rounder and their cell walls thicken as they become older. Lateral filaments with the same morphology as the initial filament are produced from the rounded cells, and grow along the surface of the substratum. Finally, upright filaments grow up into the water column bearing plurilocular or unilocular sporangia, which produce either mitospores or meiospores, respectively. Meiospores are produced through meiosis in the diploid sporophyte. Some parthenosporophytes become diploid through endoreduplication, allowing normal meiotic divisions to occur in the unilocular sporangia (Bothwell *et al*., 2010). If the parthenosporophyte remains haploid, meiosis is not possible, but unispores are produced via apomeiotic cell divisions in the unilocular sporangia (Bothwell *et al*., 2010). Therefore, both diploid and haploid *Ectocarpus* sporophytes can complete their respective sexual and asexual life cycles.

**Figure 1.**
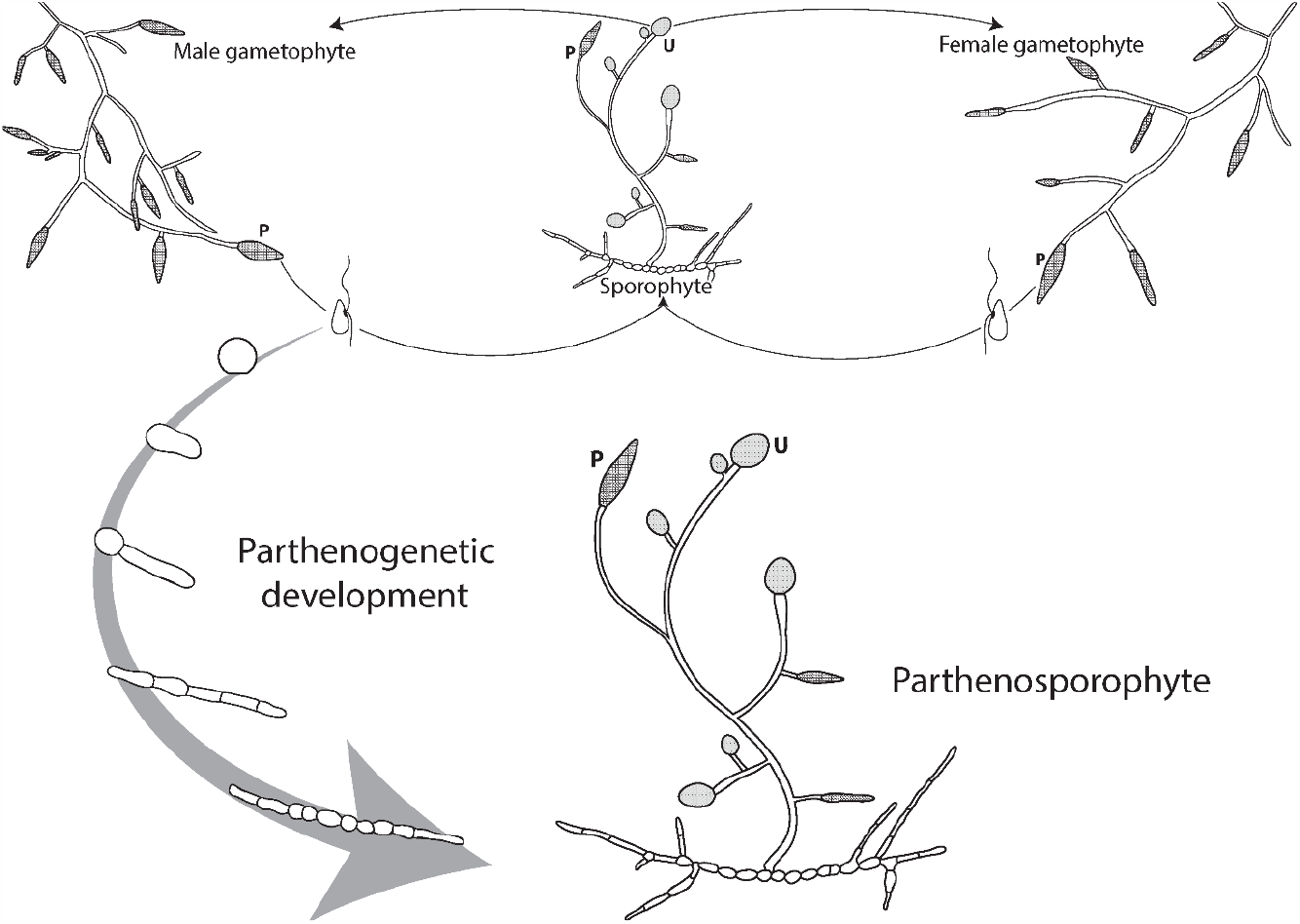
Schematic life cycle of *Ectocarpus* species 7 with emphasis on the stages relevant for parthenogenetic development. Diploid sporophytes produce haploid meiospores in unilocular sporangia (U), which develop into either male or female gametophytes. Gametophytes produce male and female gametes in plurilocular gametangia (P), which fuse to form the next diploid sporophyte generation. In the absence of fusion, gametes can develop into parthenosporophytes. After gamete settlement, the cell elongates during germination. Asynchronous bipolar germination produces two ends of a symmetric prostrate filament which grows and produces lateral filaments. Finally, upright filaments grow into the medium and form sporangia. Spores produced in unilocular sporangia of parthenoporophytes develop into gametophytes, whereas mitospores produced in the plurilocular sporangia of both sporophytes and parthenosporophytes reproduce the respective (partheno)sporophyte stage (omitted from the Figure for clarity).

While an important amount of work has been published on the early stages of development of brown algal zygotes (e.g., Brownlee and Bouget, 1998; Corellou *et al*., 2000; Bogaert *et al*., 2023), less is known about the mechanisms regulating parthenogenetic development (but see, Mignerot *et al*., 2019). The aim of our study was to characterize the dependence of the early stages of development of the *Ectocarpus* parthenosporophytes on *de novo* transcription and translation processes by applying specific inhibitors of both mechanisms. We hypothesized that triggering parthenogenesis is dependent on *de novo* transcription and translation because of the small size of the gamete, which would preclude accumulation of sufficient transcripts and proteins.

Surprisingly, we found that germination and the first cell divisions of the parthenosporophytes occurred in the presence of a transcription inhibitor, suggesting that unfertilized *Ectocarpus* gametes can initiate early development using mRNA already present in the cell. Parthenosporophytes continued to develop up to a size of 5-10 cells within 12 days, but further growth was inhibited. Germination occurred in the presence of a translation inhibitor, suggesting that the proteins necessary for germination are present in the unfertilized gamete. However, the first cell division did not occur when the translation inhibitor was present, indicating that new proteins must be produced during the first cell cycle. Our results, together with published work on the transcriptome of *Ectocarpus* gametes (Lipinska *et al*., 2013), are consistent with the view that these cells contain the mRNA and proteins necessary for the very early steps of development. As gametes are fragile and short-lived, using accumulated mRNA and proteins may be a strategy to facilitate fast embryonic development, and, in the absence of a mate, alternatively ensure asexual reproduction.

## Materials and Methods

### Algae material and culture

We used the male *Ectocarpus* species 7 (Montecinos *et al*., 2017) gametophyte strain Ec32, which has been established as a model organism and whose genome has been sequenced (Peters *et al*., 2004; Cock *et al*., 2010). Standard culture conditions were used as described by Coelho *et al*. (2012). Briefly, gametophytes were grown in 140 mm Petri dishes at 15°C, at a density of 10 individuals per petri dish. Natural sea water (NSW) was filtered, autoclaved and enriched with half-strength Provasoli nutrient solution (Provasoli-enriched seawater, PES; Starr and Zeikus, 1993). Maturity of gametophytes was accessed by microscopy. On day 0 of the experiment, all PES was removed from cultures of mature gametophytes and the material was cultivated with residual moisture in darkness at 13°C for five hours. Synchronous gamete release was then triggered by transferring the gametophytes into strong light and fresh growth medium. Inhibitors were added to the medium before gamete release to ensure that they acted on the gametes from the moment of their release from the plurilocular gametangia.

### Treatments with inhibitors

Thioluthin (Sigma-Aldrich) was stored at 1 mM in DMSO and applied at three concentrations (0.03 μM; 0.1 μM and 0.3 μM) to inhibit transcription activity by suppressing RNA polymerase function (Qiu *et al*., 2021). Emetine (Sigma-Aldrich) was stored at 1 mM in autoclaved distilled water at -20°C and was used at three concentrations (0.1 μM; 0.3 μM and 1 μM) to inhibit translation activity by irreversibly binding to the ribosomal 40S subunit, preventing polypeptide chain formation (Jiménez *et al*., 1977). The control treatment was prepared using 300 μL DMSO L^-1^ PES correspondent to the 0.3 μM thiolutin treatment. All treatments were prepared in triplicate (*n*=3). Released gametes were pipetted onto a glass coverslip inside a Petri dish and allowed to settle for 1 h before flooding the entire dish with the respective medium. The growth medium was changed every two days for a treatment duration of 12 days. Development was followed on treatment days 5, 7 and 12 by counting at least one hundred individuals under an inverted microscope. Six categories of parthenosporophyte development were quantified: round settled gametes, elongated germinated cells, two cells, 3-5 cells, 6-10 cells and larger than 10 cells. To analyze recovery from translation and transcription inhibition, inhibitors were removed by washing the parthenosporophytes three times with PES before cultivation in fresh PES for a post-culture period of 14 days.

### Microscopy

After 12 days of emetine treatment and 14 days of post-culture, emetine-treated parthenosporophytes were stained and imaged with a confocal laser scanning microscopy (SP5 TCS CLSM microscope, Leica Microsystem) equipped with a HCX PLApo 63x oil objective to visualize the effects of translation inhibition on parthenosporophyte development in comparison to parthenosporophytes grown for 26 days in control medium. Fluorescent Brightener 28 (Calcofluor white M2R, Sigma-Aldrich, excitation at 405 nm and emission at 415–475 nm) was diluted at 1 mg mL^-1^ in autoclaved distilled water, filtered at 0.2 μm and stored at -20°C. Calcofluor white stock solution was diluted 100-fold in PES, incubated with the parthenosporophytes for 15 minutes, and finally washed three times with PES before fluorescence microscopy to visualize cell walls. The plasma membranes were stained with the dye FM4-64 (ThermoFisher, T13320) at a final concentration of 5 μg mL^-1^ 30 min before imaging (excitation 561 nm, emission 575-615). The chloroplasts were visualized from the autofluorescence of the chlorophyll (excitation at 633 nm, emission 670-700 nm). All channels were acquired sequentially with a pinhole set at 1 AU with a voxel size of 150*150*378 nm (XYZ).

### Statistical analysis

To assess at which stage the development of parthenosporophytes was significantly inhibited, prevalence of developmental stages was added in descending ontogenetic order for each stage (i.e., > 10 cells, ≥ 6 cells, ≥ 3 cells, ≥ 2 cells, ≥ germinated). Linear models were fitted to assess the prevalence of developmental stages in response to inhibitor concentration within each day. Factor significance was assessed via analysis of variance (ANOVA) and *p*-values were corrected for multiple testing following the false discovery rate approach (FDR; Benjamini and Hochberg, 1995).

## Results

### Gametes exhibit limited parthenogenetic growth in the presence of a transcription inhibitor

To evaluate the role of transcription during parthenogenetic development, we monitored the effect of the transcription inhibitor thiolutin (at 0.03 μM; 0.1 μM and 0.3 μM) on the early parthenogenetic development of unfertilized *Ectocarpus* gametes. Development of the parthenosporophytes was followed for 12 days after release of the gametes (**Figure 2**). Thiolutin did not significantly affect gamete germination until day 5 (**Table 1**; **Figure 2A**; ANOVA; F_3,8_ = 2.55, *p*-value = 0.172). The first cell division was significantly delayed at day 5 only in the 0.1 μM thiolutin treatment compared to the control (ANOVA; F_3,8_ = 5.45, *p*-value = 0.049; Tukey post-hoc test, *p*-value = 0.0186). Thiolutin did not have a significant effect on the number of parthenosporophytes with ≥ 3 cells on day 5 (ANOVA; F_3,8_ = 0.14, *p*-value = 0.933). While 0.85% of control parthenosporophytes had grown to at least 6 cells on day 5, none of the thiolutin-treated parthenosporophytes had reached this stage (ANOVA; F_3,8_ = 220.11, *p*-value < 0.0001).

**Table 1.** Results of linear models assessing the effect of thiolutin on the relative prevalence of *Ectocarpus* species 7 developmental stages for each stage and time point.

|  | DFn | DFd | Day 5 |  | Day 7 |  | Day 12 |  |
| --- | --- | --- | --- | --- | --- | --- | --- | --- |
| | | | F-value | $p$-value | F-value | $p$-value | F-value | $p$-value |
| ≥ germinated | 3 | 8 | 2.5492 | 0.1719 | 0.8743 | 0.4936 | 1.000 | 0.4411 |
| $\geq 2$ cells | 3 | 8 | 5.4543 | 0.0491 | 3.518 | 0.1146 | 5.301 | 0.0440 |
| $\geq 3$-5 cells | 3 | 8 | 0.1411 | 0.9325 | 2.2772 | 0.1958 | 4.4815 | 0.0499 |
| $\geq 6$-10 cells | 3 | 8 | 220.11 | <0.0001 | 33.314 | 0.0004 | 14.803 | 0.0031 |
| > 10 cells | 3 | 8 | n/a | n/a | 3.5364 | 0.1146 | 18123 | <0.0001 |

FDR corrected $p$ -values for ANOVAs on different stages within each day. $n=3$ for each treatment.

**Figure 2.**
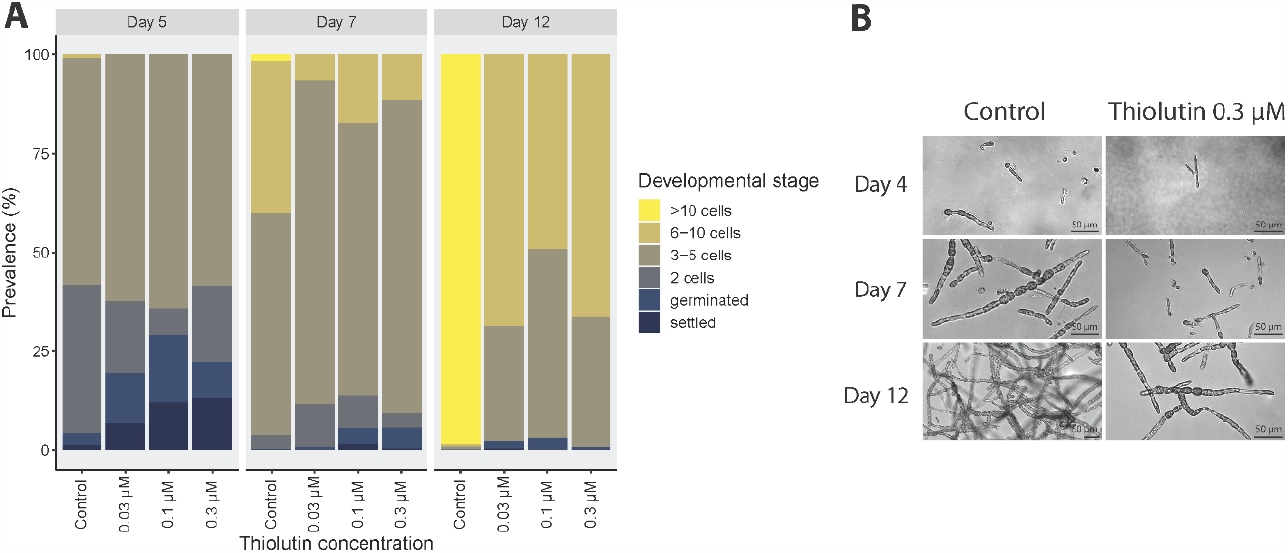
Effect of thiolutin on *Ectocarpus* species 7 parthenogenetic development. **(A)** Relative prevalence of early developmental stages of *Ectocarpus* parthenosporophytes in response to different concentrations of the transcription inhibitor thiolutin over 12 days (mean over *n*=3 per treatment and stage) compared to Provasoli-enriched natural seawater with 0.03% DMSO (control). **(B)** Microscopic documentation of the development of *Ectocarpus* parthenosporophytes over time in control medium and with 0.3 μM of thiolutin.

Continuous incubation with the inhibitor affected further cell divisions. At day 7, the proportion of parthenosporophytes with at least 6 cells was significantly lower in all thiolutin treatments (6.7–17.3%) than in the control treatment (40.1%; ANOVA; F_3,8_ = 33.31, *p*-value = 0.0004; Tukey post-hoc tests, *p*-value < 0.01). After 12 days, 98.6% of control parthenosporophytes had more than 10 cells, while thiolutin completely prevented growth beyond the 10-cell stage in all treatments (ANOVA; F_3,8_ = 18123, *p*-value < 0.0001), with 49.1–68.7% of individuals arrested at the 6-10 cell stage.

### The first cell divisions of Ectocarpus parthenosporophytes are arrested in the presence of a translation inhibitor

In addition, we investigated the effect of emetine, an inhibitor of translation, on the early development of *Ectocarpus* parthenosporophytes (**Figure 3**). Germination of parthenosporophytes was strongly affected by the 0.1 μM, 0.3 μM and 1 μM emetine treatments, as only 44.6–61.1% of the emetine-treated individuals had germinated on day 5, compared to 98.8% in the control (**Table 2; Figure 3A**; ANOVA; F_3,8_ = 87.95, *p*-value < 0.0001; Tukey post-hoc tests, *p*-value < 0.0001). In addition, the first cell divisions were strongly affected. Whereas all control individuals had undergone the first cell division after 12 days, 66.7–75.9% of emetine-treated individuals remained undivided (ANOVA; F_3,8_ = 600.3, *p*-value < 0.0001). Only 15.6–30.1% progressed to the 3-5 cell stage (ANOVA; F_3,8_ = 385.09, *p*-value < 0.0001), with a higher proportion in the 0.1 μM treatment compared to 0.3 and 1 μM (Tukey post-hoc tests, *p*-value < 0.01), indicating a dose-dependent response. Emetine fully inhibited growth beyond the 5-cell stage (**Figure 3A,B**), and induced irregular cell division planes (**Figure 3C,D**).

**Table 2.** Results of linear models assessing the effect of emetine on the prevalence of *Ectocarpus* species 7 developmental stages for each stage and time point.

|  |  |  | Day 5 |  | Day 7 |  | Day 12 |  |
| --- | --- | --- | --- | --- | --- | --- | --- | --- |
|  | DFn | DFd | F-value | p-value | F-value | p-value | F-value | p-value |
| ≥ germinated | 3 | 8 | 87.951 | <0.0001 | 17.737 | 0.0008 | 31.126 | 0.0001 |
| ≥ 2 cells | 3 | 8 | 612.46 | <0.0001 | 225.44 | <0.0001 | 600.3 | <0.0001 |
| ≥ 3-5 cells | 3 | 8 | 35.842 | 0.0001 | 562.12 | <0.0001 | 385.09 | <0.0001 |
| ≥ 6-10 cells | 3 | 8 | 220.11 | <0.0001 | 86.207 | <0.0001 | 14641 | <0.0001 |
| > 10 cells | 3 | 8 | n/a | n/a | 3.5364 | 0.0680 | 18123 | <0.0001 |

FDR corrected *p*-values for ANOVAs on different stages within each day. *n*=3 for each treatment.

**Figure 3.**
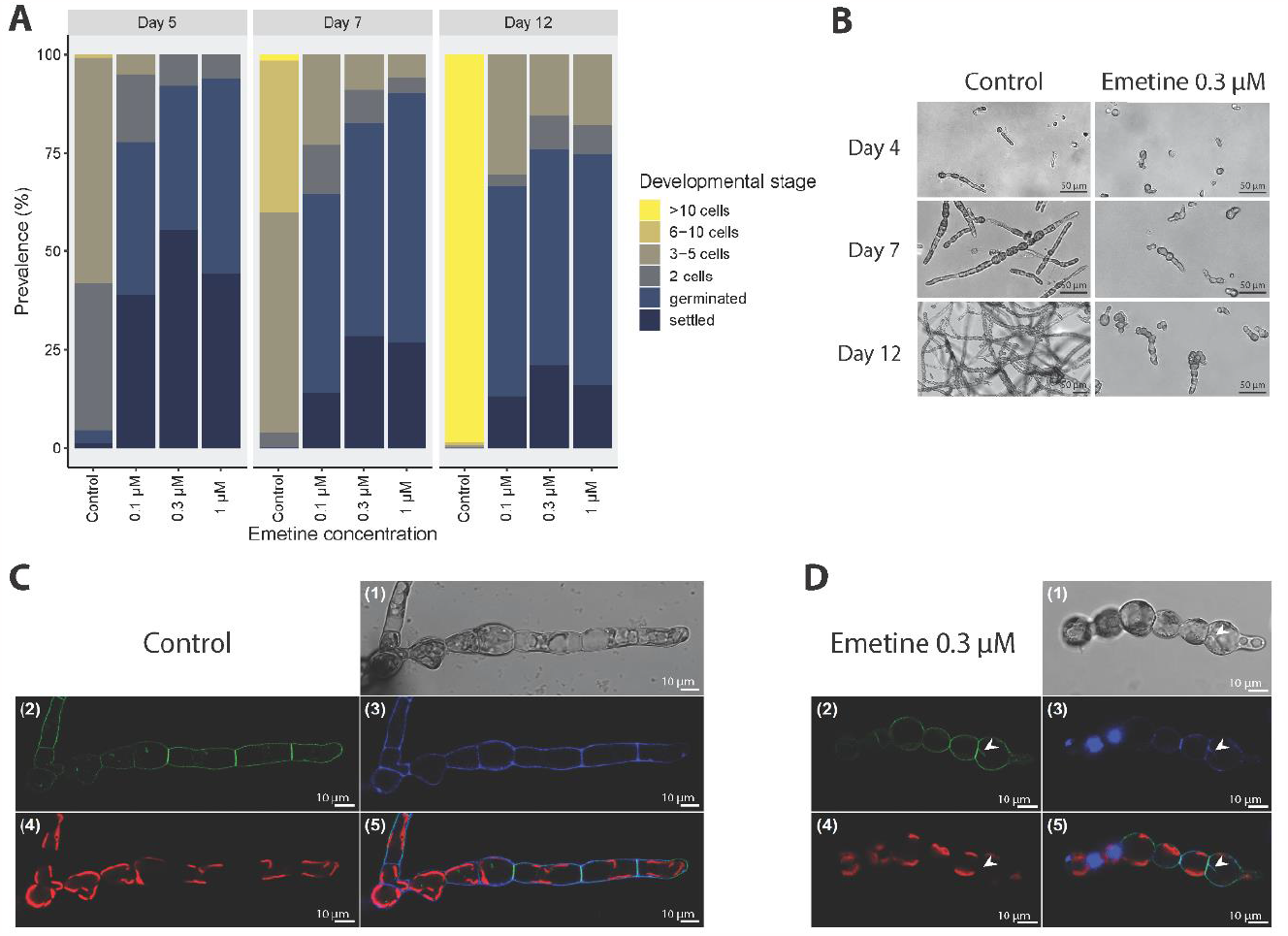
Effect of emetine on *Ectocarpus* species 7 parthenogenetic development. **(A)** Relative prevalence of early developmental stages of *Ectocarpus* parthenosporophytes in response to different concentrations of the translation inhibitor emetine over 12 days (mean over *n*=3 per treatment and stage) compared to Provasoli-enriched natural seawater with 0.03% DMSO (control). **(B)** Microphotographies (bright field) of the development of *Ectocarpus* parthenosporophytes over time in control medium and with 0.3 μM of emetine. **(C+D)** Confocal laser scanning microscopy optical sections of *Ectocarpus* parthenosporophyte filaments after **(C)** 26 days in control medium and **(D)** 12 days in 0.3 μM emetine followed by 14 days of control medium. **(1)** Bright field channel, **(2)** plasma membrane channel (FM4-64), **(3)** cell wall channel (calcofluor white M2R), **(4)** chlorophyll autofluorescence channel, **(5)** merged fluorescence channels. Arrow in **(D)** denotes an abnormal cell division plane.

### Treatment with a translation inhibitor induces persistent morphological abnormalities

Inhibition of transcription using thiolutin did not appear to result in developmental abnormalities or cell death at the tested concentrations. The treated parthenosporophytes did not exhibit unusual morphologies (**Figure 2B**) and the inhibitory effect was reversible (**Figure 4**) during the 14 days of post-culture. Thiolutin-treated parthenosporophytes recovered from the treatment and exhibited a normal, functional pattern of development at later stages, producing upright filaments and plurilocular sporangia, and showing no difference in morphology compared to control samples (**Figure 4**).

**Figure 4.**
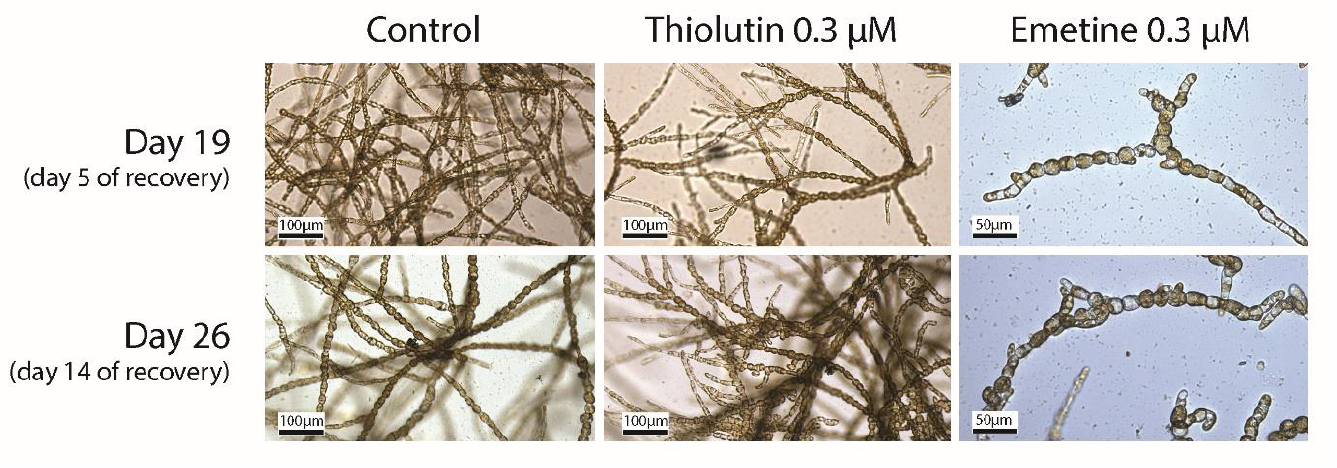
Recovery of *Ectocarpus* species 7 parthenosporophytes from inhibitor treatments. Bright field images showing *Ectocarpus* parthenosporophytes on day 19 and 26 of the experiment, corresponding to 19 and 26 days of growth in Provasoli-enriched seawater with 0.03% DMSO (control) and recovery from 12 days of treatment with 0.3 μM thiolutin or 0.3 μM emetine followed by cultivation in control medium for 5 and 14 days, respectively.

In contrast, while growth resumed during the recovery phase after 12 days of emetine treatment, the phenotypes of the resulting parthenosporophytes were abnormal even after 14 days of culture in inhibitor-free medium, including irregular cell division planes and rotund instead of elongated cell shapes (**Figures 3D; 4**). In contrast to control parthenosporophytes, which developed upright filaments after 3 weeks in culture, recovering emetine-treated parthenosporophytes did not produce upright filaments within 5 weeks.

## Discussion

### mRNA stored in the developing gamete facilitates the first parthenogenetic cell cycles

Here we investigated the early developmental pattern of *Ectocarpus* species 7 parthenosporophytes in the presence of inhibitors of transcription and translation. Our results indicated that inhibition of transcription did not affect germination of unfused male gametes and parthenogenetic development up to the 5-cell stage, while growth beyond 10 cells was fully suppressed in the presence of a transcription inhibitor (**Figure 5**). In contrast, inhibition of translation strongly affected germination and growth even beyond the treatment duration. These results suggest that early parthenogenetic development can proceed using mRNA that is already present in the *Ectocarpus* gamete when it is released from the gametangium. Germination was delayed but still took place in the presence of a translation inhibitor, suggesting that the proteins necessary for germination are already present in the unfertilized gamete. However, cell division was fully inhibited in more than 70% of individuals and no individuals grew beyond the five-cell stage, indicating that *de novo* protein synthesis is necessary for early parthenosporophyte development (**Figure 5**).

**Figure 5.**
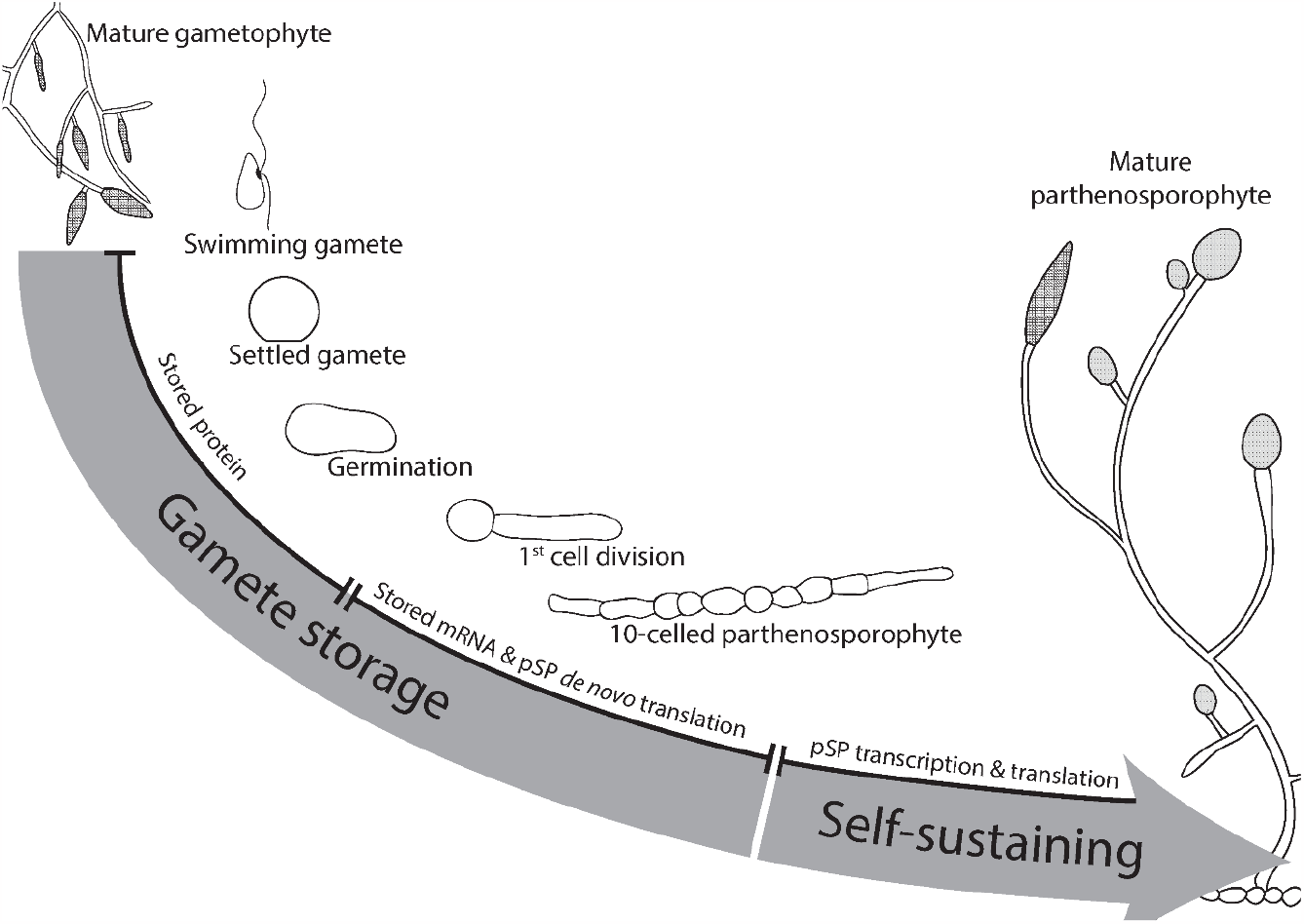
Schematic representation of contributions of mRNA and protein storage to parthenogenetic development of *Ectocarpus* species 7 gametes. Gamete settlement and germination are enabled by proteins stored in the gamete, whereas growth up to a 10-celled parthenosporophyte (pSP) is supported by the storage of mRNA, which is translated *de novo*. Beyond the 10-cell stage, the parthenosporophyte is independent of mRNA and proteins which had been stored within the developing gamete.

Previous work on the transcriptome of *Ectocarpus* gametes showed that mRNAs representing 50% of the protein coding capacity of the genome are present in the gametes despite their small size, of which 30% correspond to gamete-specific genes compared to somatic gametophyte and sporophyte tissue (Lipinska *et al*., 2013). Interestingly, enriched gene ontology (GO) terms were related to biological processes of translation, protein metabolism, and cell cycle. Furthermore, transcripts related to transcription and translation were among the 100 most abundant mRNAs in the gametes (Lipinska *et al*., 2013). mRNAs for protein metabolic processes, in particular biosynthetic pathways (ribosome and translation related) were also present in gametes. mRNAs encoding cell cycle proteins were significantly enriched in male gametes (Lipinska *et al*., 2013), including transcripts encoding mitotic kinases such as CDK1, NEK and Aurora-like kinases. Interestingly, cell cycle progression in bladder wrack (*Fucus vesiculosus*) zygotes requires maternally-inherited mRNAs coding for cyclin-dependent kinases (CDKs), which are translated only after fertilization (Corellou *et al*., 2001). Additional highly expressed genes in *Ectocarpus* gametes include cyclins involved in the G1/S and G2/M transitions of the cell cycle (cyclinD3, cyclin A, cyclin B), and Smc4, a subunit of condensin, which is involved in chromosome assembly and segregation during mitosis, and which is a substrate of CDK1 in yeast (Ubersax *et al*., 2003). According to our data, parthenosporophytes can proceed through five successive cell divisions in the presence of a transcription inhibitor, suggesting that mRNAs present in the gametes are sufficient to support growth, cell division and metabolic processes necessary for the first steps of development of the *Ectocarpus* parthenosporophyte. In contrast to cell cycle genes, genes related to photosynthesis were underrepresented in male gametes (Lipinska *et al*., 2013), indicating that the demand for photosynthetic energy in developing gametes is low, which may be compensated by the presence of storage lipids in *Ectocarpus* gametes (Maier, 1997).

Interestingly, male parthenogenesis is not universal in the genus *Ectocarpus*. For instance, male gametes of some strains of the closely related species *Ectocarpus siliculosus* are not capable of developing parthenogenetically beyond the first few cell divisions, whereas female gametes can develop into functional parthenosporophytes (Mignerot *et al*., 2019). In the study carried out by Mignerot *et al*. (2019), three quantitative trait loci located at the sex locus and on an autosome were identified as controlling parthenogenesis. The arrested development of non-parthenogenetic *E. siliculosus* male gametes is reminiscent of that of thiolutin-treated *E*. species 7 (strain Ec32) parthenosporophytes in our study, in that development of the *E. siliculosus* germlings arrests at the 3-5 cell stage and results in death after 20 days (Mignerot *et al*., 2019). Therefore, it may be possible that the male gametes of some strains of *E. siliculosus* are not able to complete parthenogenetic development because they lack the storage of essential mRNA and proteins.

### Early developmental processes are essential for the normal patterning of the adult alga

Continuous incubation of *Ectocarpus* parthenosporophytes with emetine inhibited the first cell division. Following the removal of the inhibitor, cell divisions were re-initiated. However, the development of parthenosporophytes was deeply affected, indicating that early developmental processes are essential for the normal patterning of later stage *Ectocarpus* parthenosporophytes. In many multicellular organisms, early stages of zygote development are critical for the determination of different cell fates of the early embryonic cells (Fucus: Bouget *et al*., 1998; Brownlee and Bouget, 1998; *C. elegans*: Schneider and Bowerman, 2003; *Arabidopsis*: ten Hove *et al*., 2015). Our data suggest that developmental events in the initial cell impact embryonic development in *Ectocarpus*, affecting the morphology of newly formed cells and preventing the growth of upright filaments. It remains unclear whether these long-lasting developmental defects were directly caused by the disruption of crucial developmental processes during the emetine treatment, or by the disturbance of the first cell division as a second-order effect.

Two three amino acid loop extension homeodomain transcription factors (TALE HD TFs) have been identified as central to the control of the sporophyte generation developmental program in *Ectocarpus* (*ORO* and *SAM*; Coelho *et al*., 2011; Arun *et al*., 2019). Similarly as described for two TALE HD TFs in *Chlamydomonas*, representing the lineage of green algae and land plants (Lee *et al*., 2008), they are able to heterodimerize, which is presumed to have an important function in triggering the sporophyte developmental program (Arun *et al*., 2019). Whereas in *Chlamydomonas*, either TALE HD TF is exclusively expressed in gametes of only one sex, transcripts of both *ORO* and *SAM* are highly abundant in both sexes of *Ectocarpus* gametes (Arun *et al*., 2019), but it is not yet known whether they are translated. In the moss *Physcomitrella patens*, four life-cycle related TALE HD TF proteins are present in eggs, but at least one of them (BELL1) cannot be detected in male gametangia or sperm (Horst *et al*., 2016). BELL1 has a crucial role in zygote development, it accumulates to high amounts in the embryo (Horst *et al*., 2016), and its transcription is controlled by a glutamate receptor (GLR2; Ortiz-Ramírez *et al*., 2017). Speculatively, *Ectocarpus* TALE HD TF translation or heterodimerization may therefore be activated by an upstream regulatory pathway, which is triggered upon gamete fusion, but delayed in unfertilized gametes.

### Parthenogenesis as a potentially adaptive trait?

*Ectocarpus* is a sessile broadcast spawner, synchronously releasing gametes into open water, many of which may not fuse with a partner of the opposite sex (Müller, 1966). Furthermore, the lack of cell walls in gametes directly exposes them to their abiotic environment and facilitates grazing, which makes them the one of the most vulnerable stages of the *Ectocarpus* life cycle (Coelho *et al*., 2000; Roleda *et al*., 2005; Müller *et al*., 2008). It may therefore be an adaptive strategy to provide gametes with the necessary cellular machinery and substrates to initiate development, rather than relying on the availability of sufficient photosynthetic energy and nutrients for *de novo* transcription and translation. Following gamete fusion, this may facilitate rapid development into a more resilient and self-sustaining sporophyte which completes the sexual life cycle. However, even when sexual reproduction fails, some *Ectocarpus* gametes are able to develop into parthenosporophytes which circumvents the loss of unfused propagules and may ensure vegetative reproduction. On the contrary, no parthenotes were identified among hundreds of *Ectocarpus* individuals from NW France and SW Italy (Couceiro *et al*., 2015), indicating that if parthenogenesis does occur in wild populations, the fitness of parthenosporophytes may be lower than that of sexually obtained sporophytes in the same environment. However, fully parthenogenetic populations of the brown alga *Scytosiphon lomentaria* (Ectocarpales) have been described in Japan (Hoshino *et al*., 2019), which potentially inhabit ecological niches unsuitable for sexual strains (Hoshino *et al*., 2021).

In conclusion, our results are consistent with the idea that the mRNA stored in developing male *Ectocarpus* sp. 7 gametes at the time of release is sufficient to initiate parthenogenetic growth, but that development rapidly requires *de novo* translation, as the protein stock of gametes is only sufficient to support gamete germination and, to a lesser extent, early development up to the 5-cell stage. Considering that *Ectocarpus* sp. 7 is a sessile broadcast spawner with both asexual and sexual life cycles, the mRNA pool in *Ectocarpus* gametes may be adapted to allow fast propagation and population maintenance even under conditions that are suboptimal for sexual reproduction, by ensuring an alternative vegetative pathway which allows clonal reproduction until mates are available again.

## Acknowledgements

We thank Audrina Plaisance for help with the *Ectocarpus* cultures and counting.

## Author contributions

RL performed the experiments, DL analyzed the data, SC performed the fluorescence microscopy, JM was involved in study conception and data analysis, JMC and SMC supervised the project, DL and RL wrote the manuscript which was revised and approved by all authors.

## References

Arun A, Coelho SM, Peters AF, et al. 2019. Convergent recruitment of TALE homeodomain life cycle regulators to direct sporophyte development in land plants and brown algae. eLife 8: e43101.

Benjamini Y, Hochberg Y. 1995. Controlling the false discovery rate: A practical and powerful approach to multiple testing. Journal of the Royal Statistical Society: Series B (Methodological) 57: 289–300.

Bogaert KA, Arun A, Coelho SM, De Clerck O. 2013. Brown algae as a model for plant organogenesis In: De Smet I, ed. Methods in Molecular Biology. Plant Organogenesis. Totowa, NJ: Humana Press, 97–125.

Bogaert KA, Zakka EE, Coelho SM, De Clerck O. 2023. Polarization of brown algal zygotes. Seminars in Cell & Developmental Biology 134: 90–102.

Bothwell JH, Marie D, Peters AF, Cock JM, Coelho SM. 2010. Role of endoreduplication and apomeiosis during parthenogenetic reproduction in the model brown alga Ectocarpus. New Phytologist 188: 111–121.

Bouget F-Y, Berger F, Brownlee C. 1998. Position dependent control of cell fate in the Fucus embryo: role of intercellular communication. Development 125: 1999–2008.

Brownlee C, Bouget F-Y. 1998. Polarity determination in Fucus: From zygote to multicellular embryo. Seminars in Cell & Developmental Biology 9: 179–185.

Brownlee C, Bouget F-Y, Corellou F. 2001. Choosing sides: Establishment of polarity in zygotes of fucoid algae. Seminars in Cell & Developmental Biology 12: 345–351.

Cock JM, Sterck L, Rouzé P, et al. 2010. The Ectocarpus genome and the independent evolution of multicellularity in brown algae. Nature 465: 617–621.

Coelho SM, Cock JM. 2020. Brown algal model organisms. Annual Review of Genetics 54: 71–92.

Coelho SM, Godfroy O, Arun A, Le Corguillé G, Peters AF, Cock JM. 2011. OUROBOROS is a master regulator of the gametophyte to sporophyte life cycle transition in the brown alga Ectocarpus. Proceedings of the National Academy of Sciences 108: 11518–11523.

Coelho SM, Peters AF, Müller D, Cock JM. 2020. Ectocarpus: an evo-devo model for the brown algae. EvoDevo 11: 19.

Coelho SM, Rijstenbil JW, Brown MT. 2000. Impacts of anthropogenic stresses on the early development stages of seaweeds. Journal of Aquatic Ecosystem Stress and Recovery 7: 317–333.

Coelho SM, Scornet D, Rousvoal S, et al. 2012. How to cultivate Ectocarpus. Cold Spring Harbor Protocols 2012: pdb.prot067934.

Corellou F, Brownlee C, Detivaud L, Kloareg B, Bouget FY. 2001. Cell cycle in the fucus zygote parallels a somatic cell cycle but displays a unique translational regulation of cyclindependent kinases. The Plant Cell 13: 585–98.

Corellou F, Coelho SMB, Bouget F-Y, Brownlee C. 2005. Spatial re-organisation of cortical microtubules in vivo during polarisation and asymmetric division of Fucus zygotes. Journal of Cell Science 118: 2723–2734.

Corellou F, Potin P, Brownlee C, Kloareg B, Bouget F-Y. 2000. Inhibition of the establishment of zygotic polarity by protein tyrosine kinase inhibitors leads to an alteration of embryo pattern in Fucus. Developmental Biology 219: 165–182.

Couceiro L, Le Gac M, Hunsperger HM, et al. 2015. Evolution and maintenance of haploiddiploid life cycles in natural populations: The case of the marine brown alga Ectocarpus. Evolution 69: 1808–1822.

om Dieck I. 1992. North Pacific and North Atlantic digitate Laminaria species (Phaeophyta): hybridization experiments and temperature responses. Phycologia 31: 147–163.

Heesch S, Serrano-Serrano M, Barrera-Redondo J, et al. 2021. Evolution of life cycles and reproductive traits: Insights from the brown algae. Journal of Evolutionary Biology 34: 992–1009.

Horst NA, Katz A, Pereman I, Decker EL, Ohad N, Reski R. 2016. A single homeobox gene triggers phase transition, embryogenesis and asexual reproduction. Nature Plants 2: 15209.

Hoshino M, Hiruta SF, Croce ME, et al. 2021. Geographical parthenogenesis in the brown alga Scytosiphon lomentaria (Scytosiphonaceae): Sexuals in warm waters and parthenogens in cold waters. Molecular Ecology 30: 5814–5830.

Hoshino M, Okino T, Kogame K. 2019. Parthenogenetic female populations in the brown alga Scytosiphon lomentaria (Scytosiphonaceae, Ectocarpales): Decay of a sexual trait and acquisition of asexual traits (M Cock, Ed.). Journal of Phycology 55: 204–213.

en Hove CA, Lu K-J, Weijers D. 2015. Building a plant: Cell fate specification in the early Arabidopsis embryo. Development 142: 420–430.

Jiménez A, Carrasco L, Vazquez D. 1977. Enzymic and nonenzymic translocation by yeast polysomes. Site of action of a number of inhibitors. Biochemistry 16: 4727–4730.

Lee J-H, Lin H, Joo S, Goodenough U. 2008. Early sexual origins of homeoprotein heterodimerization and evolution of the plant KNOX/BELL family. Cell 133: 829–840.

Lipinska AP, D’hondt S, Van Damme EJ, De Clerck O. 2013. Uncovering the genetic basis for early isogamete differentiation: A case study of Ectocarpus siliculosus. BMC Genomics 14: 909.

Luthringer R, Cormier A, Ahmed S, Peters AF, Cock JM, Coelho SM. 2014. Sexual dimorphism in the brown algae. Perspectives in Phycology 1: 11–25.

Maier I. 1997. The fine structure of the male gamete of Ectocarpus siliculosus (Ectocarpales, Phaeophyceae). I. General structure of the cell. European Journal of Phycology 32: 241–253.

Mignerot L, Avia K, Luthringer R, et al. 2019. A key role for sex chromosomes in the regulation of parthenogenesis in the brown alga Ectocarpus (J de Meaux, Ed.). PLOS Genetics 15: e1008211.

Montecinos AE, Couceiro L, Peters AF, Desrut A, Valero M, Guillemin M-L. 2017. Species delimitation and phylogeographic analyses in the Ectocarpus subgroup siliculosi (Ectocarpales, Phaeophyceae) (M Cock, Ed.). Journal of Phycology 53: 17–31.

Müller DG. 1966. Untersuchungen zur Entwicklungsgeschichte der Braunalge Ectocarpus siliculosus aus Neapel. Planta 68: 57–68.

Müller DG. 1967. Generationswechsel, Kernphasenwechsel und Sexualität der Braunalge Ectocarpus siliculosus im Kulturversuch. Planta 75: 39–54.

Müller DG, Murúa P, Westermeier R. 2019. Reproductive strategies of Lessonia berteroana (Laminariales, Phaeophyceae) gametophytes from Chile: Apogamy, parthenogenesis and cross-fertility with L. spicata. Journal of Applied Phycology 31: 1475–1481.

Müller R, Wiencke C, Bischof K. 2008. Interactive effects of UV radiation and temperature on microstages of Laminariales (Phaeophyceae) from the Arctic and North Sea. Climate Research 37: 203–213.

Ortiz-Ramírez C, Michard E, Simon AA, et al. 2017. GLUTAMATE RECEPTOR-LIKE channels are essential for chemotaxis and reproduction in mosses. Nature 549: 91–95.

Peters AF, Marie D, Scornet D, Kloareg B, Cock JM. 2004. Proposal of Ectocarpus siliculosus (Ectocarpales, Phaeophyceae) as a model organism for brown algal genetics and genomics. Journal of Phycology 40: 1079–1088.

Peters AF, Scornet D, Ratin M, et al. 2008. Life-cycle-generation-specific developmental processes are modified in the immediate upright mutant of the brown alga Ectocarpus siliculosus. Development 135: 1503–1512.

Pool JE, Vejlupkova Z, Goodner BW, Lu G, Quatrano RS. 2004. Localization to the rhizoid tip implicates a Fucus distichus Rho family GTPase in a conserved cell polarity pathway. Planta 219.

Qiu C, Malik I, Arora P, et al. 2021. Thiolutin is a direct inhibitor of RNA Polymerase II. bioRxiv: doi: 10.1101/2021.05.05.442806.

Robinson KR, Miller BJ. 1997. The Coupling of cyclic GMP and photopolarization of Pelvetia zygotes. Developmental Biology 187: 125–130.

Roleda MY, Wiencke C, Hanelt D, Van De Poll WH, Gruber A. 2005. Sensitivity of Laminariales zoospores from Helgoland (North Sea) to ultraviolet and photosynthetically active radiation: implications for depth distribution and seasonal reproduction. Plant, Cell and Environment 28: 466–479.

Schneider SQ, Bowerman B. 2003. Cell polarity and the cytoskeleton in the Caenorhabditis elegans zygote. Annual Review of Genetics 37: 221–249.

Silberfeld T, Leigh JW, Verbruggen H, Cruaud C, de Reviers B, Rousseau F. 2010. A multi-locus time-calibrated phylogeny of the brown algae (Heterokonta, Ochrophyta, Phaeophyceae): Investigating the evolutionary nature of the “brown algal crown radiation”. Molecular phylogenetics and evolution 56: 659–74.

Starr RC, Zeikus JA. 1993. UTEX—The culture collection of algae at the University of Texas at Austin. 1993 list of cultures. Journal of Phycology 29: 1–106.

Ubersax JA, Woodbury EL, Quang PN, et al. 2003. Targets of the cyclin-dependent kinase Cdk1. Nature 425: 859–864.

